# Identification of the first oomycete mating-type locus sequence in the grapevine downy mildew pathogen, *Plasmopara viticola*

**DOI:** 10.1101/2020.02.26.962936

**Authors:** Yann Dussert, Ludovic Legrand, Isabelle D. Mazet, Carole Couture, Marie-Christine Piron, Rémy-Félix Serre, Olivier Bouchez, Pere Mestre, Silvia Laura Toffolatti, Tatiana Giraud, François Delmotte

## Abstract

Mating types are self-incompatibility systems that promote outcrossing in plants, fungi and oomycetes. Mating-type genes have been widely studied in plants and fungi, but have yet to be identified in oomycetes, eukaryotic organisms closely related to brown algae that cause many destructive animal and plant diseases. We identified the mating-type locus of *Plasmopara viticola*, the oomycete responsible for grapevine downy mildew, one of the most damaging grapevine diseases worldwide. Using a genome-wide association approach, we identified a 570 kb repeat-rich non-recombining region controlling mating types, with two highly divergent alleles. We showed that one mating type was homozygous, whereas the other was heterozygous at this locus. The mating-type locus encompassed 40 genes, including one encoding a putative hormone receptor. Our findings have fundamental implications for our understanding of the evolution of mating types, as they reveal a unique determinism involving an asymmetry of heterozygosity, as in sex chromosomes and unlike other mating-type systems. This identification of the mating-type locus in such an economically important crop pathogen also has applied implications, as outcrossing facilitates rapid evolution and resistance to harsh environmental conditions.

## INTRODUCTION

Mating systems control the degree of outcrossing in natural populations, thus affecting adaptability (Lande & Schemske 1985; Charlesworth & Charlesworth 1987; Charlesworth *et al.* 1990; Igic *et al.* 2008; Lande & Porcher 2015). Indeed, outcrossing promotes gene flow and, therefore, the rapid spread of beneficial alleles, the generation of new allelic combinations and the purging of deleterious alleles. Various genetic systems have evolved across the tree of life to enforce outcrossing, such as separate sexes or mating types (Beukeboom & Perrin 2014). The sex of many plants and animals is determined by sex chromosomes (Beukeboom & Perrin 2014). Mating types, preventing mating between individuals carrying the same alleles, have evolved independently in many lineages of the tree of life, including fungi, ciliates, green algae and oomycetes (Billiard *et al.* 2012; Sekimoto 2017; Orias *et al.* 2017). Plants possess a molecular system called self-incompatibility, which enforces outcrossing and is analogous to mating type (Fujii *et al.* 2016). Given the fundamental importance of mating types in life cycles and evolution, their molecular determinism has been extensively studied, particularly in plants, fungi and ciliates. Mating types are controlled by a number of different mechanisms, even within these groups (Billiard *et al.* 2012; Fujii *et al.* 2016; Orias *et al.* 2017).

Mating type systems enforcing outcrossing at the diploid stage, involving only two mating types, have been described in oomycetes on the basis of cross incompatibilities (Judelson 2009). However, no genomic sequence for a mating-type locus has yet been identified in this lineage. Oomycetes are diploid eukaryotic organisms closely related to diatoms and brown algae (Lévesque 2011). This group includes a number of animal and plant pathogens causing significant environmental and economic damage. Oomycetes are responsible for damaging diseases, such as saprolegniosis, a lethal disease affecting wild and farmed fish (van West 2006), sudden oak death and downy mildew in several crops (e.g., *Bremia lactucae*, responsible for lettuce downy mildew, and *Plasmopara halstedii*, responsible for sunflower downy mildew), and the infamous potato blight caused by *Phytophthora infestans*, responsible for the Irish potato famine in the 1840s.

Mating types are of the utmost importance in these organisms, as they control gamete compatibility, and sexual reproduction generates thick-walled spores called oospores that can resist harsh conditions and survive for several years. Outcrossing also produces recombinant genotypes, which can mediate faster adaptation to control measures. For example, invasive populations of *Ph. infestans* were not able to reproduce sexually for a long period, due to the lack of one mating type, but the recent introduction of the missing mating type has resulted in higher genetic diversity in some areas (Brurberg *et al.* 2011), and the emergence of an aggressive lineage (Gavino *et al.* 2000). In oomycetes, the mating-type locus has been located only on a genetic linkage map in *Phytophthora* species (Judelson *et al.* 1995; Fabritius & Judelson 1997; Van Der Lee *et al.* 2004) and *Bremia lactucae* (Sicard *et al.* 2003), and the only known mating-type factors are purified hormones in *Phytophthora spp.* (Qi *et al.* 2005; Harutyunyan *et al.* 2008; Ojika *et al.* 2011). However, no genes determining mating type have yet been identified in oomycetes.

*Plasmopara viticola*, a pathogen causing grapevine downy mildew, one of the most devastating grapevine diseases worldwide, is an oomycete. *Plasmopara viticola* was introduced into European vineyards from North America in the 1870s (reviewed in Fontaine *et al.* 2013) and rapidly spread, invading vineyards all over the world. The life cycle of *Pl. viticola* includes an asexual multiplication phase during the spring and summer and a sexual reproduction event in the fall, generating the thick-walled sexual spores (oospores) required for overwintering (Vercesi *et al.* 2010). In *Pl. viticola*, only two mating types have been observed, and mating can occur only between diploid individuals of different mating types (Wong *et al.* 2001). This results in high rates of outcrossing, a key element explaining rapid adaptation to fungicides (Toffolatti *et al.* 2011; Delmas *et al.* 2017) and to resistant cultivars (Peressotti *et al.* 2010; Delmas *et al.* 2016) in this species. We identified the first oomycete mating-type locus sequence, with an approach combining phenotypic analysis (mating types determined by crosses) with genome-wide single-nucleotide polymorphisms (SNPs) in *Pl. viticola*.

## RESULTS

We determined the mating type of 54 diploid individuals of *Pl. viticola* collected across Europe (Supplementary Table 1), in experimental crosses against six testers, three for each of the two mating types, arbitrarily called P1 and P2. The P1 mating type was inferred for an individual if it produced oospores when inoculated on grapevine leaves with P2 testers but not with P1 testers, and conversely for P2. We identified 26 individuals of the P1 mating type and 28 of the P2 mating type (Supplementary Table 2). We sequenced the genomes of these 54 diploid individuals with short-read technology, and mapped the reads onto a recent high-quality reference sequence obtained by long-read sequencing with high coverage (Dussert *et al.* 2019). We retained 2.011 million SNPs after filtering. No genetic subdivision associated with mating types was detected in population structure analyses based on principal component analysis or clustering analysis applied to a dataset filtered for SNPs in close linkage disequilibrium (LD) (Supplementary Figure 1). Conditions were therefore favorable for genetic-phenotype association studies.

**Table 1.**
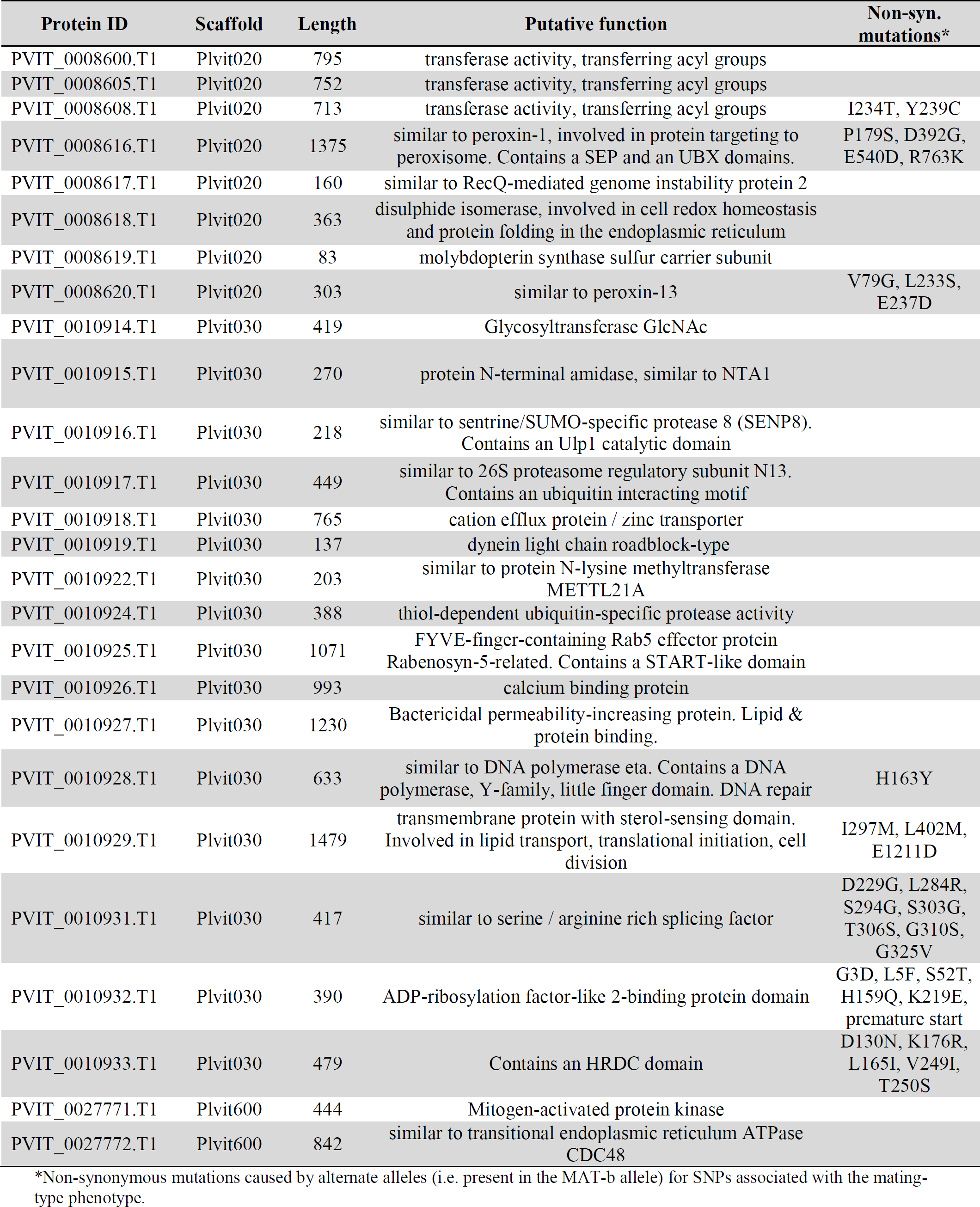
Putative function of genes in the mating-type locus of *Plasmopara viticola*.

**Figure 1.**
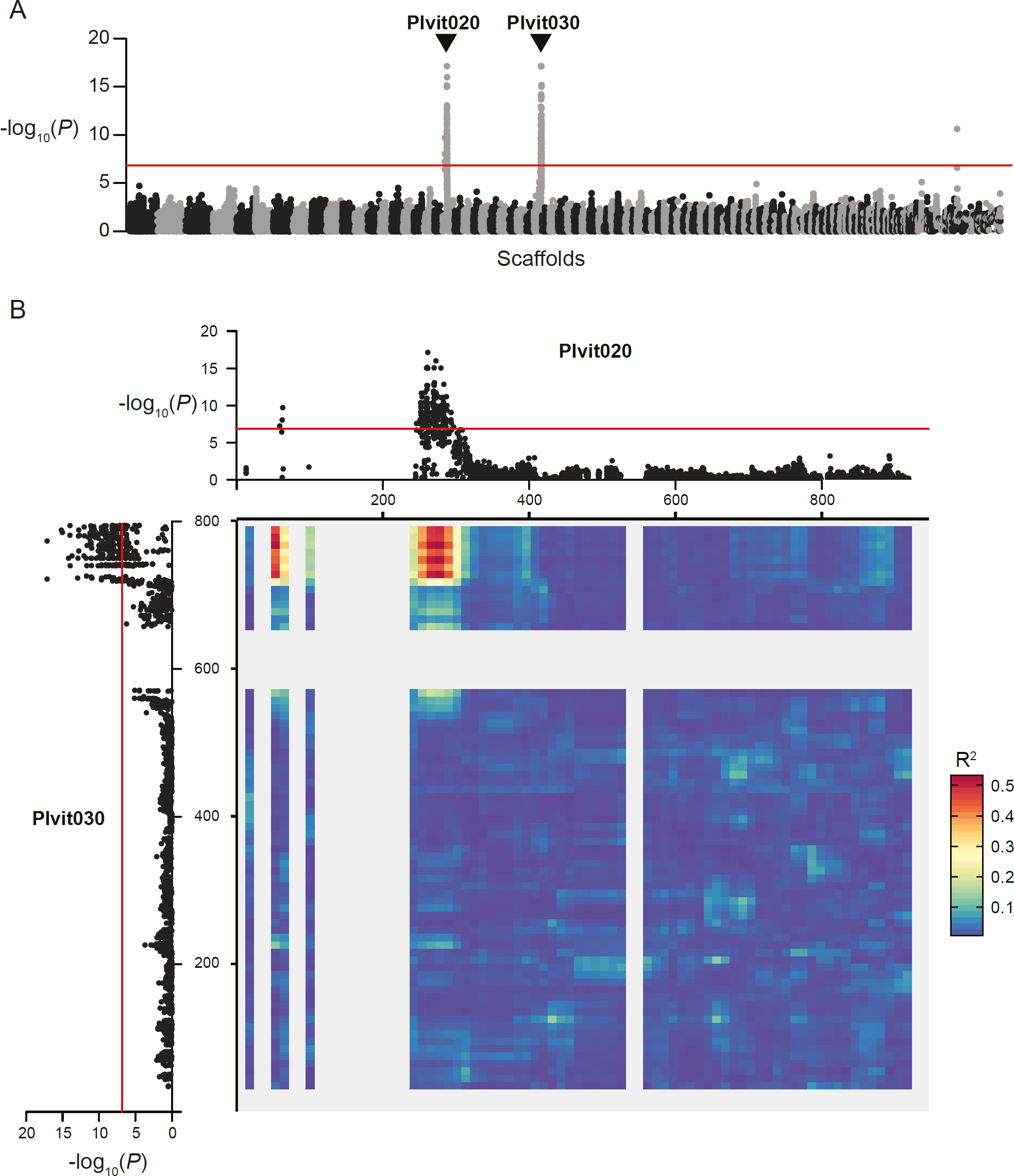
Genome-wide association analysis for identifying mating-type regions in *Plasmopara viticola*. A: Manhattan plot of the negative log_10_-transformed association *P*-values between mating type and SNPs along the *Pl. viticola* genome. Alternating black and grey blocks of dots mark the limits between scaffolds. The two scaffolds with a significant association signal (Plvit020 and Plvit030) are indicated with arrows. Only 5% of the SNPs with a *P*-value < 0.1 are represented, to keep the number of plotted points manageable. B: Manhattan plots of the negative log10-transformed association *P*-values between mating type and SNPs for Plvit020 and Plvit030, and linkage disequilibrium between the two scaffolds represented as a heatmap (80 bins). The significance threshold for the association analysis, computed with 10,000 permutations, is represented as a red line in both panels.

Using a genome-wide association approach, we identified two genomic regions with significant signals of association with the mating-type phenotype, located at the edges of the scaffolds Plvit020 and Plvit030 (Figure 1). SNPs at these two scaffold extremities were in very strong LD (Figure 1B), indicating that they were located in the same genomic region and that this region probably lacked recombination. The rate of LD decay was much lower in this region than in the rest of the genome (Supplementary Figure 2), providing further support for the hypothesis of a lack of recombination. The incomplete assembly of this locus in the reference genome was probably due to its high repeat content. Indeed, we observed two large regions composed exclusively of tandem repeat arrays covering 327 kb (Supplementary Figure 3).

**Figure 2.**
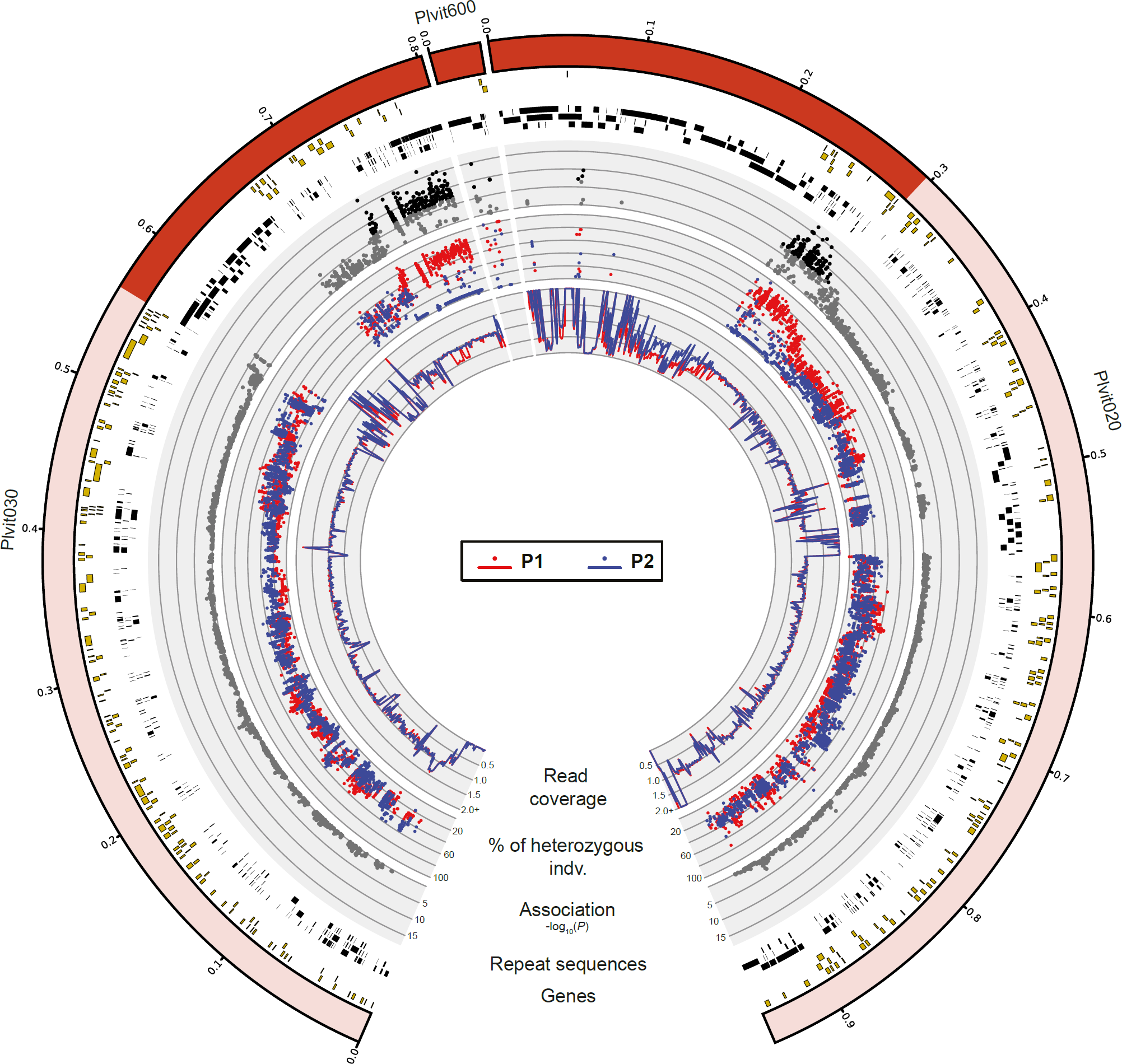
Genomic regions associated with mating type in *Plasmopara viticola*. The regions significantly associated with the mating-type phenotype are shown in dark red within the scaffold ideograms (outer track). Annotated genes are represented as yellow rectangles, and repeated sequences as black rectangles. Significant negative log_10_-transformed *P*-values from the association analysis are represented by black dots, and other values are represented by gray dots. The percentage of heterozygous individuals for each SNP is shown for individuals of mating types P1 by red dots and P2 by blue dots. The mean normalized read coverage (for adjacent 1 kb windows) is shown by red lines for P1 individuals and blue lines for P2 individuals. The data were obtained with the reference-based SNP calling approach for the scaffolds Plvit020 and Plvit030 and the reference-free SNP calling approach for Plvit600.

**Figure 3.**
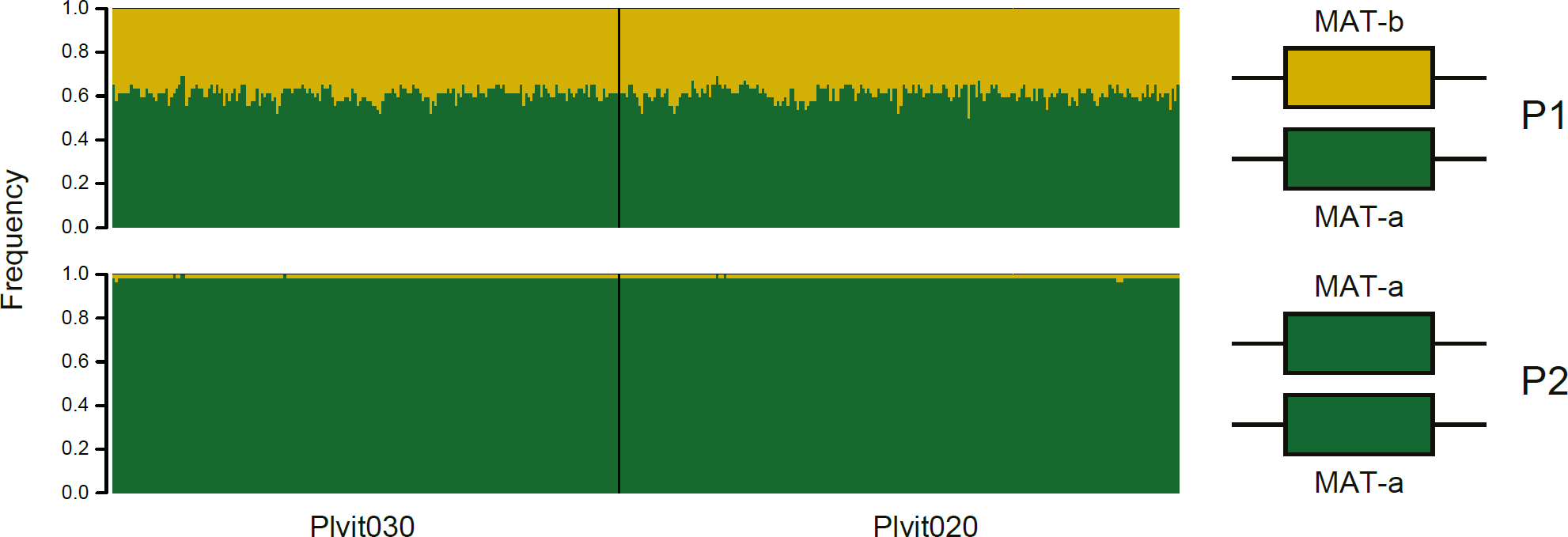
Mating type is determined by a two-allele system in *Plasmopara viticola*. Left panel: Allele frequencies at SNPs associated with the mating-type phenotype in P1 individuals (top) and P2 individuals (bottom). Only results for the reference-based SNP calling approach are shown. Right panel: Proposed model for mating-type determination, with homozygosity in P2 individuals and heterozygosity in P1 individuals. For both panels, the reference allele (i.e. the allele found in the reference genome) is shown in green, and the alternative allele is shown in yellow.

We therefore used a reference-free SNP calling method to identify potential missing sequences between the two scaffolds. We detected short genomic sequences carrying SNPs with significant signals of association with the mating-type phenotype that were missing from the reference assembly (see Material & Methods section). We reassembled the PacBio reads corresponding to these short sequences and obtained a new contig, Plvit600 (32,220 bp), with SNPs significantly associated with the mating-type phenotype in strong linkage disequilibrium with SNPs located at the edges of Plvit020 and Plvit030, and therefore probably located between these two scaffolds (Supplementary Figure 4). Plvit600 also included a large region entirely composed of tandem repeat arrays (Supplementary Figure 3). We therefore concluded that the mating-type locus was at least 570 kb long (Figure 2). P2 individuals were homozygous for the reference allele (designated MAT-a) at the mating-type locus, whereas P1 individuals were heterozygous, carrying the MAT-a reference allele and a second allele, called MAT-b (Figure 3). We found no homozygous individual for the alternative MAT-b allele. This suggested that the MAT-b allele is dominant. The two alleles were highly differentiated along the 570 kb, as shown by the large difference of heterozygosity levels (Figure 2), again consistent with a lack of recombination in this region.

**Figure 4.**
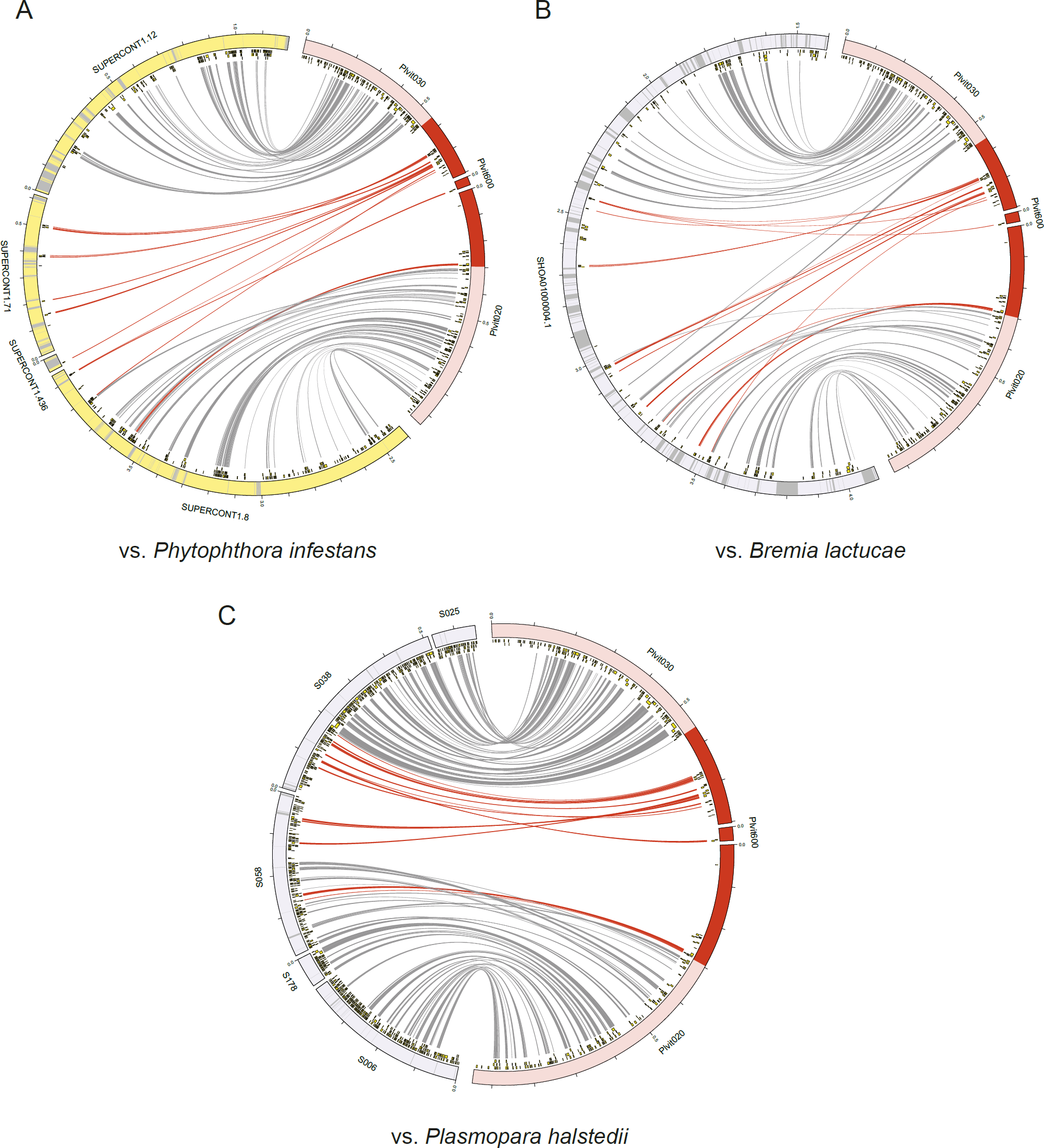
Orthologs of the *Plasmopara viticola* mating-type locus genes in the genomes of *Phytophthora infestans* (A), *Bremia lactucae* (B) and *Plasmopara halstedii* (C). Orthology relationships were determined with reciprocal best hits and are represented by links between ideograms. The regions significantly associated with the mating-type phenotype in *Pl. viticola* and the links corresponding to genes in these regions are shown in dark red. Predicted genes are represented by yellow boxes, and the grayed-out regions in ideograms represent gaps in the assemblies.

The mating-type region included a total of 40 predicted coding sequences (Supplementary Table 3), 26 of which had predicted functions and did not correspond to TEs (Table 1). Based on the predicted functions of these genes, the most promising candidate for involvement in mating-type determination was a gene encoding a transmembrane protein (PVIT_0010929.T1) with a sterol-sensing domain and lipid transport activity. This protein might act as a hormone receptor, and hormones have been identified as mating-type factors during initiation of sexual reproduction in *Phytophthora spp.* (Qi *et al.* 2005; Harutyunyan *et al.* 2008; Ojika *et al.* 2011). Another two genes (PVIT_0010925.T1 and PVIT_0010927.T1) encoded proteins with lipid-binding domains that could potentially interact with mating hormones. We also found a putative mitogen-activated protein (MAP) kinase gene (PVIT_0027771.T1) in the identified mating-type region. MAP kinases play an important role in the mating pheromone response pathway in fungi (Zhao *et al.* 2007), and genes encoding kinases are present at the mating-type loci of some fungi (Karos *et al.* 2000). We also found genes encoding a protein with an HRCD (helicase and RNaseD C-terminal; PVIT_0010933.T1) domain and a RecQ-mediated genome instability protein 2 (PVIT_0008617.T1). Proteins of this type can interact with DNA and are often associated with RecQ helicases and, thus, DNA repair. These proteins may also be involved in recombination suppression or gene regulation. This is potentially also the case for a Cdc48-like protein (PVIT_0027772.T1) encoded by a gene present at the mating-type locus. Cdc48 can interact with protein complexes on chromatin and is involved in gene regulation and cell cycle progression. Finally, we also found four genes (PVIT_0008616.T1, PVIT_0010916.T1, PVIT_0010917.T1 and PVIT_0010924.T1) encoding proteins with ubiquitin-like or ubiquitin-interacting domains, which could potentially be involved in addressing proteins to the proteasome and, thus, in protein degradation.

Using a reciprocal best hit approach with annotated whole proteomes, we identified the mating-type region in the genomes of *Ph. infestans*, and two other species more closely related to *Pl. viticola*: *Br. lactucae* and *Pl. halstedii*. The genes orthologous to putative mating-type genes were found on three scaffolds for *Ph. infestans*, one for *Br. lactucae* and two for *Pl. halstedii* (Figure 4, Supplementary Table 4). The genomic region including the candidate genes differed in size between species, probably reflecting the difference in repeat content between the species genomes (Haas *et al.* 2009; Pecrix *et al.* 2019; Dussert *et al.* 2019). We observed differences of gene order between species, suggesting the existence of genomic rearrangements of the mating-type locus between species, as expected for non-recombining mating-type regions (Badouin *et al.* 2015).

## DISCUSSION

We report here the first identification of an oomycete mating-type locus sequence, in the genome of *Pl. viticola*. The *Pl. viticola* mating-type locus was found in a 570 kb non-recombining, repeat-rich region of the genome encompassing 40 genes. The region was heterozygous (MAT-a/MAT-b) in the P1 mating type and homozygous (MAT-a/MAT-a) in the P2 mating type, indicating dominance of the MAT-b allele. These findings are consistent with previous findings for marker segregation analyses in *Phytophthora* species and *Br. lactucae* (Judelson 1996; Fabritius & Judelson 1997; Sicard *et al.* 2003). This genetic determinism of mating type is unique in the tree of life and resembles determinism based on sex chromosomes, with one sex typically being homozygous and the other heterozygous. By contrast, fungal mating types are determined at the haploid stage, with mating occurring between gametes carrying different mating-type alleles, and many species harboring multiple alleles. Diploid or dikaryotic fungal individuals are necessarily heterozygous at the mating type locus in heterothallic species and can thus undergo selfing (Giraud *et al.* 2008). Mating types in fungi therefore do not prevent diploid selfing, they only prevent same-clone mating (Billiard *et al.* 2012), while mating types in oomycetes and plants do prevent diploid selfing. Self-incompatibility systems in plants are similar to oomycete mating types, as they prevent same-allele mating, and also display multiple alleles (Fujii *et al.* 2016).

As expected for non-recombining regions under long-term balancing selection, we found the two *Pl. viticola* mating types to be highly divergent. The mating-type locus in *Ph. infestans* is also heteromorphic, with highly differentiated alleles, hemizygous fragments and genomic translocations, and it is associated with deleterious recessive alleles (Judelson 1996; Randall *et al.* 2003; Van Der Lee *et al.* 2004). The high degree of divergence between alleles, the number of rearrangements, the high repeat content, and the sheltering of deleterious alleles by permanent heterozygosity are typical of sex and mating-type chromosomes in plants, animals and fungi (Bachtrog 2005; Graves 2006; Badouin *et al.* 2015; Charlesworth 2015).

The localization of the mating-type genomic sequence and elucidation of the determinism of mating type (homozygous versus heterozygous) in an oomycete, and the identification of orthologous mating-type loci in other oomycete species will have fundamental implications for our understanding of the evolution of mating types in this lineage, and should shed light on why several other oomycetes have no self-incompatibility systems and can self, a state known as homothallism in oomycetes (Judelson 2009). In fungi, homothallic species often have the two mating-type alleles in each haploid genome (Butler 2014). Our study also paves the way for further investigations of the proximal molecular mechanisms determining mating type in oomycetes. We identified possible mating type-determining genes, including one encoding a putative hormone receptor. This may guide the development of innovative control methods based on disruption of the sexual cycle of the pathogen, since infections from oospores play a major role during grapevine downy mildew epidemics (Gobbin *et al.* 2005). Furthermore, the identification of the mating-type locus in such an economically important crop pathogen may improve our understanding of pathogen adaptation, as sex and outcrossing promote rapid evolution and resistance to harsh conditions.

## MATERIALS AND METHODS

### Sampling and preparation of the study material

Grapevine leaves infected with *Pl. viticola* were collected across Europe (Supplementary Table 1). Diploid oomycete individuals were isolated from diseased leaves by dilution to isolate single spores, as follows: isolates of *Pl. viticola* were first propagated on detached leaves of V*itis vinifera* cv. Cabernet-Sauvignon. Sporangia were collected and suspended in water at a concentration of 10^4^ sporangia/mL (estimated using Malassez cells) and serial dilutions were performed to obtain a concentration of 0.1 sporangia/10 µL. We applied 10 µL of this inoculum to leaf disks, which were then placed in a growth chamber at 22°C (12 h light/12 h dark photoperiod) for seven days. We retained a single sporulating disk per isolate, which was, thus, derived from a single diploid sporangium.

### DNA extraction and whole-genome resequencing

Detached leaves from *V. vinifera* cv. Cabernet-Sauvignon or Muscat Ottonel plants were sterilized with bleach, washed with sterilized water and infected with fresh *Pl. viticola* inoculum. Leaves were placed in a growth chamber for 4-7 days at 21-22°C (12 h/12 h or 16 h/8 h light/dark photoperiod). Sporangiophores and sporangia were then collected in distilled water. The suspension was centrifuged at maximum speed, the supernatant was removed and the pellet was stored at −80 °C. DNA was extracted with a modified CTAB procedure (Dussert *et al.* 2019) or with the Qiagen DNeasy Plant Mini Kit.

Genomic DNA was sequenced by Beckman Coulter Genomics (Grenoble, France) or at the GeT-PlaGe facility (Toulouse, France), on an Illumina HiSeq 2000 sequencer (2×100 bp paired-end reads) or an Illumina HiSeq 3000 sequencer (2×150 bp). Read lengths and mean genome coverage are provided in Supplementary Table 1.

### Mating-type determination

Six *Pl. viticola* individuals (Supplementary Tables 1 & 2) were chosen as mating-type testers: three individuals of each mating type. These testers were first crossed with each other in all possible pairings, for separation of the six individuals into two opposite mating-type groups, P1 and P2 (three individuals each). We then crossed 54 individuals with these six testers to determine their mating type. In all experiments, leaf disks from *V. vinifera* cv. Cabernet-Sauvignon plants were placed on 20 g/L agar in Petri dishes and co-inoculated with 15 µL droplets containing 4×10^4^ sporangia/mL from each individual (three droplets per disk, six replicate disks per pairing). After four to five days in a growth chamber at 22°C (12 h light/12 h dark photoperiod), disks were examined for sporulation, to check for successful inoculation. The leaf disks were then transferred to a growth chamber at 12°C (12 h light/12 h dark photoperiod) for about three weeks. Each leaf disk was then observed with a binocular microscope, to check for the presence of oospores, indicative of successful mating.

### Read mapping and single-nucleotide polymorphism (SNP) calling

Reads from each individual were mapped onto the *Pl. viticola* PacBio reference genome (Dussert *et al.* 2019) with the Glint aligner v1.0.rc12.826:833 (available at https://forge-dga.jouy.inra.fr/projects/glint), allowing a maximum of 15% mismatches, to handle genomic regions with high heterozygosity (other parameters: mappe mode, --no-lc-filtering --best-score -- lrmin 80), and filtering out reads with a mapping quality below 3 and reads that were not properly paired.

A pileup file was generated from the resulting alignment files with SAMtools 1.3.1 (Li *et al.* 2009) mpileup without probabilistic alignment quality computation (-B parameter). SNPs and short insertions and deletions (indels) were then called with the multi-sample procedure (mpileup2snp and mpileup2indel) of VARSCAN 2.4.3 (Koboldt *et al.* 2012) (--min-coverage 10 - -min-reads2 5 --min-avg-qual 15 --min-var-freq 0.2 --min-freq-for-hom 0.75 --p-value 0.001 -- strand-filter 1). For each individual, sites with a read coverage more than 1.5 standard deviations on either side of the mean value for genome coverage were discarded with BCFtools 1.1-60-g3d5d3d9 (Li 2011). SNPs located within 2 bp of a detected indel were also filtered out. Finally, only sites with less than 10% missing data were retained. After filtering, the dataset contained 2,011 million SNPs.

### Genetic diversity and association of SNPs with mating types

We used SAMtools to assess the normalized read coverage for 1 kb windows for each individual. Using only SNPs with a minor allele frequency (MAF) below 0.1, the percentage of heterozygous individuals for each site along the genome was calculated separately for P1 and P2 individuals with VCFtools (Danecek *et al.* 2011). Linkage disequilibrium between sites, as assessed with the R^2^ statistic, was also computed with VCFtools. Finally, linkage disequilibrium decay along scaffolds was analyzed with PopLDdecay 3.28 (Zhang *et al.* 2019).

SNP thinning was performed by clumping (as described in Privé *et al.* 2018) with PLINK 1.07 (Purcell *et al.* 2007), with MAF as the statistic of importance and a LD threshold (R^2^) of 0.2. Population genetic structure was investigated by principal component analysis (PCA) with PCAdapt (Luu *et al.* 2017) in R (R Core Team 2017). We also investigated population clustering by computing ancestry coefficients for individuals (Q-matrix) with the non-negative matrix factorization algorithm sNMF (Frichot *et al.* 2014) implemented in the LEA package (Frichot & François 2015) in R. We set the number of groups *K* between 1 and 10, with 50 runs for each value of *K*. The optimal number of groups was chosen using the minimum cross-entropy criterion.

For identification of the genomic region controlling mating types, we analyzed the association between genotype and mating type (genome-wide association approach) with TASSEL 5.2.41 (Bradbury *et al.* 2007) on SNPs with no missing data and a MAF above 0.1, using a GLM model with the Q-matrix estimated by sNMF. *P*-values were corrected with 10,000 permutations (threshold: 0.05). We filtered out the small number of association signals found in short scaffolds, corresponding to isolated SNPs in repeat regions.

### Identification of genomic regions associated with mating type and absent from the reference genome

The discovery of SNPs associated with mating type in the edges of two scaffolds suggested that the mating-type locus had been only partially assembled in the reference assembly.

We therefore used discoSnp++ (Peterlongo *et al.* 2017), a reference-free method for identifying SNPs. Briefly, SNPs are detected from bubbles found in a de Bruijn graph built from raw sequencing reads, and short contigs are reconstructed around the SNPs. This analysis was carried out with a subset of 30 individuals (Supplementary Table 1), as computing resource issues arose if the analysis was extended to a larger number of individuals. We used a *k*-mer size of 31, the smart branching strategy (-b 1), a maximum of five SNPs in unique bubbles (-P 5), and polymorphisms were extended with left and right contigs (-T). Only sites with less than 10% missing data and a MAF below 0.1 were retained. Contigs were mapped onto the reference genome with the VCF_creator module of discoSnp++ and an association analysis was carried out with TASSEL, as described above, to check that our results were robust to the SNP calling method.

Contigs from discoSnp++ including SNPs significantly associated with mating type were aligned with the self-corrected PacBio reads used for reference genome assembly (Dussert *et al.* 2019) with BLASTN+, retaining hits displaying at least 95% identity and 95% query coverage. PacBio reads corresponding to these hits were assembled with SMARTdenovo (available from https://github.com/ruanjue/smartdenovo) using default parameters. Assembled contigs were aligned against the reference scaffolds including SNPs with a significant association signal, with NUCmer from the MUMmer 3.23 package (Kurtz *et al.* 2004) using default parameters, and only regions absent from the reference assembly were retained. Available paired-end reads and 3 kb mate-pair reads (Dussert *et al.* 2016) and the self-corrected PacBio reads were aligned against the reference genome and the new mating-type contigs, with BWA 0.7.12 (Li & Durbin 2009) for short reads and NGMLR (Sedlazeck *et al.* 2018) for long reads. Aligned reads were used to polish the new mating-type contigs with Pilon 1.22 (Walker *et al.* 2014).

After sequence polishing, contigs from the discoSNP analysis were again mapped, but this time to the reference assembly and the newly assembled contigs, and a final association analysis was carried out. Newly assembled contigs with no SNP significantly associated with the matin type were discarded. For the remaining contigs, repeat sequences were annotated with TEannot from REPET 2.2 (Quesneville *et al.* 2005; Flutre *et al.* 2011), using the *Pl. viticola* repeat library (Dussert *et al.* 2019). Available RNA-seq data (Mestre *et al.* 2016; Dussert *et al.* 2019) were aligned against the reference assembly and the new contigs with STAR 2.4 (Dobin *et al.* 2013), and were used as hints for gene prediction in the new contigs with AUGUSTUS 3.1 (Stanke *et al.* 2008), together with the training files produced by the previous annotation (Dussert *et al.* 2019). Proteins for which functional domains of transposable elements were detected with InterProScan 5 (Jones *et al.* 2014) or for which hits were obtained with TransposonPSI (http://transposonpsi.sourceforge.net; 30% coverage, e-value < 1e^-10^) were filtered out. The remaining proteins were aligned against the NCBI nr database with BLASTP+ (-evalue 1e^-5^ - max_hsp 20 -num_alignment 20), and functionally annotated with Blast2GO 4.1.9 (Gotz *et al.* 2008) using the BLASTP+ and InterProScan results for gene ontology (GO) term mapping.

For all candidate genes within the mating-type locus (i.e. from the scaffolds in the reference assembly and the new contigs), we further checked whether coding sequences corresponded to transposable elements: protein sequences for those genes were masked with CENSOR 4.2.29 (Kohany *et al.* 2006) using RepBase 23.08, which allowed us to compute for each protein a percentage of coverage from known transposable elements.

### Comparative genomics with *Phytophthora infestans, Bremia lactucae* and *Plasmopara halstedii*

We identified the orthologs of the mating-type candidate genes in the annotated proteomes of *Ph. infestans* (Haas *et al.* 2009, ASM14294v1 assembly), *Br. lactucae* (Fletcher *et al.* 2019) and *Pl. halstedii* (Pecrix *et al.* 2019), with the reciprocal best hits (RBH) method using BLASTP+ (-evalue 1e^-6^ -qcov_hsp_perc 50 –use_sw_tback, min. identity 40%). Results were visualized with Circos (Krzywinski *et al.* 2009).

## DATA AVAIBILITY

Raw reads for each individual will be deposited in GenBank (see Supplementary Table 1 for SRA accession numbers) and analysis files have been deposited in Dataverse (doi.org/10.15454/ILQ12T).

## ACKNOWLEDGMENTS

We thank Vincent Thomas for helping with the production of material and Pal Kozma (University of Pécs, Hungary), Mauro Jermini (Agroscope, Switzerland), Sara Legler (UCSC, Italy) Hervé Steva (Bordeaux, France) and Atanas Atanassov (JGC, Bulgaria) for providing *Pl. viticola* isolates. We are grateful to Jérôme Gouzy (INRAE, Toulouse) for his help and suggestions concerning data analysis. We thank the Genotoul bioinformatics platform Toulouse Midi-Pyrenees (Bioinfo Genotoul) for providing help and computing resources. This work was performed in collaboration with the GeT core facility, Toulouse, France (http://get.genotoul.fr). This study was supported by the European Commission (INNOVINE, FP7-KBBE-311775), and the French National Research Agency (GANDALF project, ANR-12-ADAP-0009; EFFECTOORES project, ANR-13-ADAP-0003; Investments for the future program in the Cluster of Excellence COTE, ANR-10-LABX-45; Investments for the future program France Génomique National infrastructure, ANR-10-INBS-09). TG acknowledges receipt of the ERC advanced grant EvolSexChrom #832252.

