## Supplementary Material for "Identification of the first oomycete mating-type locus sequence in the grapevine downy mildew pathogen, *Plasmopara viticola*"

[1]INRAE, Bordeaux Sciences Agro, Université Bordeaux, SAVE, F-33140, Villenave d'Ornon, France; [2]LIPM, INRAE, Université de Toulouse, CNRS, Castanet-Tolosan, France; [3] Université de Strasbourg, INRAE, SVQV UMR-A 1131, F-68000 Colmar, France; [4]INRAE, US 1426, GeT-PlaGe, Genotoul, Castanet-Tolosan, France; [5]Dipartimento di Scienze Agrarie e Ambientali, Università degli Studi di Milano, Milano, Italy; [6]Ecologie Systematique et Evolution, CNRS, AgroParisTech, Universite Paris-Saclay, 91400 Orsay France

[†] Current affiliation: School of Biological and Chemical Sciences, Queen Mary University of London, London, UK

**This document includes:**

Supplementary Figure 1 Population structure analysis in *Plasmopara viticola*, based on a LD-thinned dataset of 35,516 SNPs.

Supplementary Figure 2 Linkage disequilibrium (LD) decay in *Plasmopara viticola*.

Supplementary Figure 3 Tandem repeat arrays in the region associated with the mating type phenotype in *Plasmopara viticola*.

Supplementary Figure 4 Genome-wide association analysis for identifying mating-type regions in *Plasmopara viticola*, based on a reference-free SNP calling method.

Supplementary Table 1 *Plasmopara viticola* individuals used for the mating type study.

Supplementary Table 2 Mating-type phenotyping for *Plasmopara viticola* individuals.

Supplementary Table 3 Functional annotation of all candidate genes in the mating-type locus of *Plasmopara viticola*.

Supplementary Table 4 Orthologs of the *Plasmopara viticola* mating type genes in the genomes of *Phytophthora infestans*, *Bremia lactucae* and *Plasmopara halstedii*.

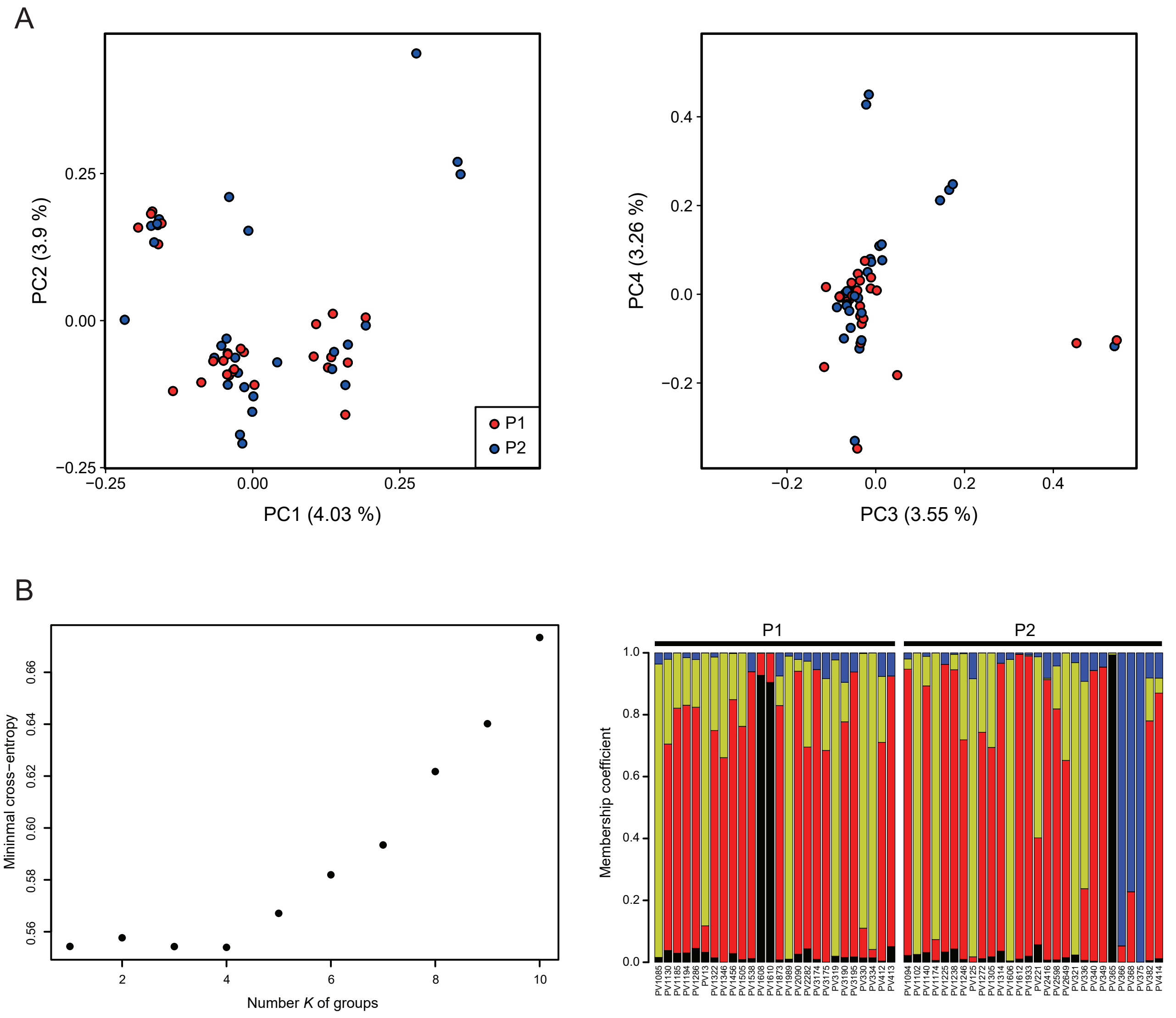


**Supplementary Figure 1 Population structure analysis in *Plasmopara viticola*, based on a LD-thinned dataset of 35,516 SNPs.** A: Principal component analysis. Individuals with the P1 mating type are represented in red and individuals with the P2 mating type in blue. The percentage of variation explained by each principal component is indicated in brackets. B: Population clustering by the non-negative matrix factorization algorithm sNMF. The values of minimum cross-entropy for each number *K* of groups are represented in the left panel. The membership coefficients for each individual for the run with the lower (i.e. better) minimum cross-entropy at *K* = 4 are represented in the right panel. Individuals are represented by vertical lines, divided into four colored sections corresponding to the individuals' membership coefficient in each group.

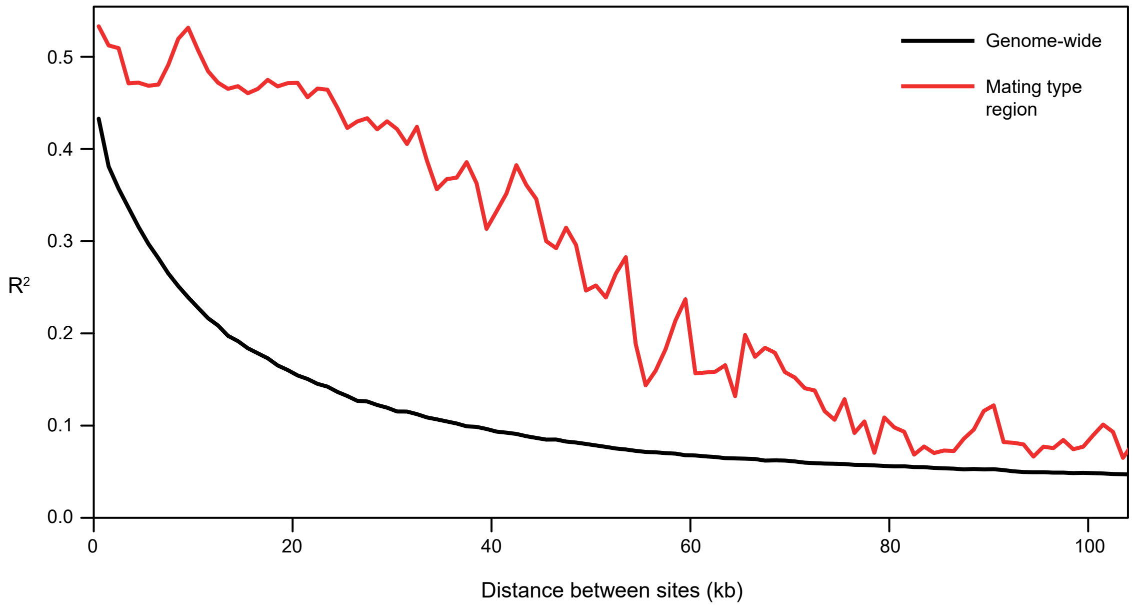


**Supplementary Figure 2 Linkage disequilibrium (LD) decay in *Plasmopara viticola*.** The LD between SNPs, measured by $R^2$, is represented as a function of physical distance between SNPs, with $R^2$ values averaged for windows of 1000 bp. The LD decay curve is represented in black for the whole genome, and in red for the region associated with the mating-type phenotype in the Plvit020 and Plvit030 scaffolds.

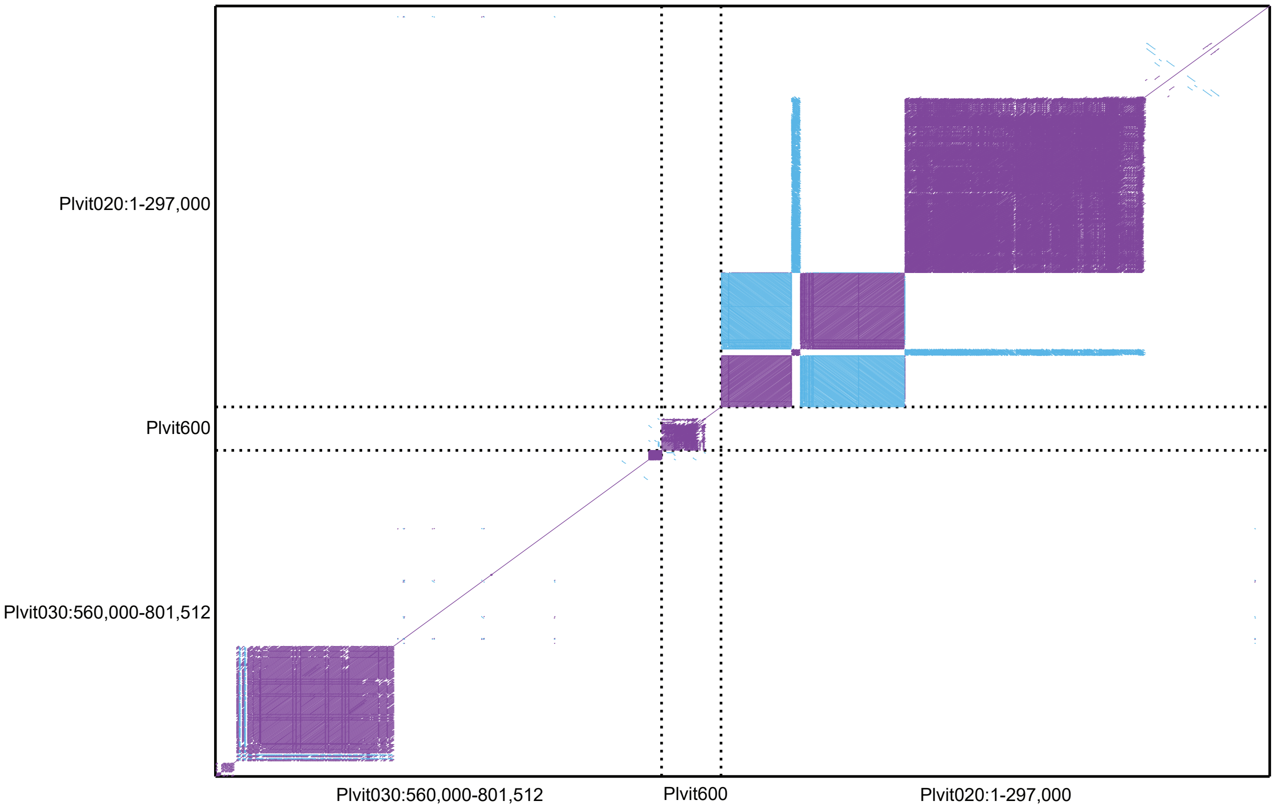


**Supplementary Figure 3 Tandem repeat arrays in the region associated with the mating type phenotype in *Plasmopara viticola*.** All-versus-all alignments for sequences associated with the mating type are represented as a dot plot, with forward matches in purple and reverse matches in blue. Results are shown for the two scaffolds found in the reference genome (Plvit020 and Plvit030) and a third scaffold (Plvit600) assembled using data from the reference-free SNP calling method (see main text).

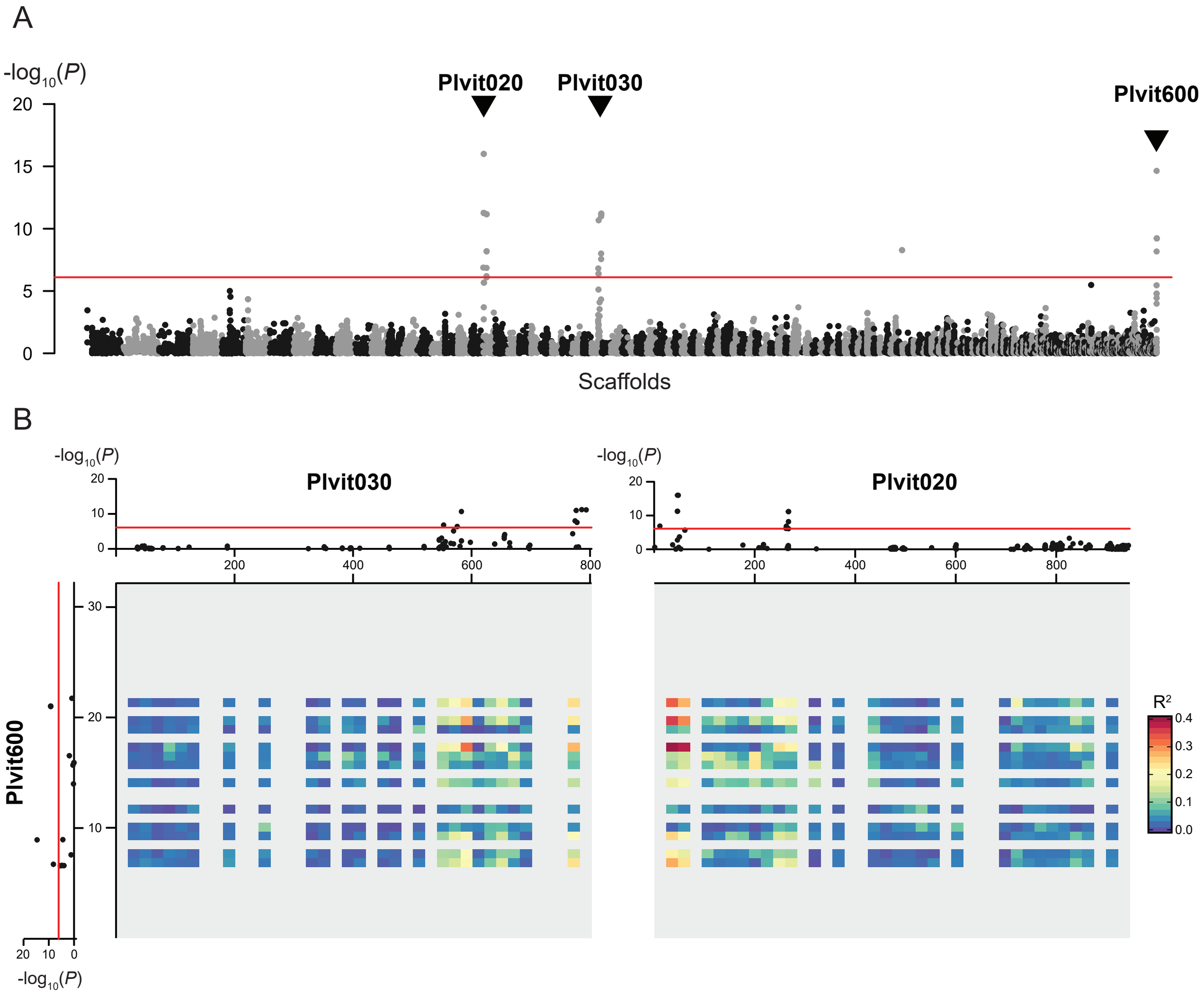


**Supplementary Figure 4 Genome-wide association analysis for identifying mating-type regions in *Plasmopara viticola*, based on a reference-free SNP calling method.** A: Manhattan plot of the negative $log_{10}$-transformed association *P*-values between mating type and SNPs along the *Pl. viticola* genome. Alternating black and grey blocks of dots mark the limits between scaffolds. Scaffolds with a significant association signal (Plvit020, Plvit030 and Plvit600) are indicated with arrows. Plvit600 was assembled using data from the reference-free SNP calling method (see main text). B: Manhattan plots of the negative $log_{10}$-transformed association *P*-values between mating type and SNPs for Plvit020, Plvit030 and Plvit600, and linkage disequilibrium between Plvit600 and the two other scaffolds, represented as a heatmap (40 bins). The significance threshold for the association analysis, computed with 10,000 permutations, is represented as a red line in both panels.

**Supplementary Table 1** ***Plasmopara viticola*** **individuals used for the mating type study**

| ID | Country | Location | Latitude (decimal degrees) | Longitude (decimal degrees) | Read length (bp) | Mean coverage | Used for reference-free SNP analysis | SRA accessions |
|---|---|---|---|---|---|---|---|---|
| PV13 | France | Latresne | 44.79 | -0.50 | 100 | 57.1 | | |
| PV125 | Hungary | Pècs | 46.08 | 18.23 | 100 | 107.3 | | |
| PV221* | France | Blanquefort | 44.92 | -0.62 | 100 | 100.7 | X | SRX1970160, SRX1970161, SRX1970162 |
| PV319 | France | Côte d'Or | NA | NA | 100 | 93.7 | X | |
| PV321 | Germany | Kröv | 49.99 | 7.09 | 100 | 144.1 | X | |
| PV330 | Germany | Pfaffenweiler | 47.94 | 7.76 | 100 | 60.3 | | |
| PV334 | Germany | Ehrenkirchen | 47.91 | 7.75 | 100 | 79.2 | X | |
| PV336 | Hungary | Eger | 47.91 | 20.38 | 100 | 47.0 | X | |
| PV340 | Hungary | Tolcsva | 48.28 | 21.44 | 150 | 23.0 | | |
| PV349 | Switzerland | Cugnasco | 46.18 | 8.92 | 150 | 19.4 | X | |
| PV365 | Switzerland | Cugnasco | 46.18 | 8.92 | 100 | 45.0 | | |
| PV366 | Switzerland | Cugnasco | 46.18 | 8.92 | 150 | 26.3 | | |
| PV368 | Switzerland | Cugnasco | 46.18 | 8.92 | 150 | 19.3 | X | |
| PV375 | Switzerland | Cugnasco | 46.18 | 8.92 | 100 | 149.7 | | |
| PV382 | Switzerland | Cugnasco | 46.18 | 8.92 | 150 | 33.8 | | |
| PV412 | Switzerland | Cugnasco | 46.18 | 8.92 | 150 | 37.5 | | |
| PV413* | Switzerland | Cugnasco | 46.18 | 8.92 | 150 | 23.5 | X | |
| PV414 | Switzerland | Cugnasco | 46.18 | 8.92 | 150 | 27.8 | | |
| PV1085 | Switzerland | Chexbres | 46.48 | 6.77 | 100 | 126.8 | X | |
| PV1094 | Switzerland | Chexbres | 46.48 | 6.77 | 150 | 24.0 | X | |
| PV1102 | Switzerland | Nyon | 46.40 | 6.23 | 100 | 115.2 | | |
| PV1130 | Switzerland | Leytron | 46.19 | 7.22 | 150 | 32.7 | | |
| PV1140 | Switzerland | Leytron | 46.19 | 7.22 | 150 | 23.3 | X | |
| PV1174 | Germany | Ihringen | 48.05 | 7.62 | 100 | 168.0 | X | |
| PV1185 | Germany | Ihringen | 48.05 | 7.62 | 150 | 30.0 | | |
| PV1194 | Germany | Ihringen | 48.05 | 7.63 | 150 | 28.8 | X | |
| PV1225 | Germany | Ihringen | 48.05 | 7.63 | 150 | 18.7 | X | |
| PV1238* | Germany | Pfaffenweiler | 47.94 | 7.76 | 150 | 23.0 | X | |
| PV1246 | Germany | Ehrenkirchen | 47.93 | 7.73 | 150 | 36.4 | | |
| PV1272 | Germany | Ihringen | 48.05 | 7.62 | 150 | 32.3 | X | |
| PV1286 | Germany | Ehrenkirchen | 47.93 | 7.73 | 150 | 27.5 | X | |
| PV1305 | Germany | Ebringen | 47.96 | 7.78 | 150 | 33.8 | | |
| PV1314 | Germany | Ebringen | 47.96 | 7.78 | 150 | 17.9 | | |
| PV1322 | Germany | Ehrenkirchen | 47.93 | 7.73 | 150 | 29.6 | | |
| PV1346 | Germany | Pfaffenweiler | 47.93 | 7.75 | 150 | 34.8 | | |
| PV1400* | Germany | Pfaffenweiler | 47.94 | 7.75 | - † | - † | | |
| PV1456 | Germany | Pfaffenweiler | 47.94 | 7.76 | 150 | 28.4 | X | |
| PV1505 | Germany | Pfaffenweiler | 47.94 | 7.76 | 150 | 30.9 | | |
| PV1538* | Czech Republic | NA | NA | NA | 150 | 21.7 | X | |
| PV1606 | Switzerland | Leytron | 46.19 | 7.22 | 100 | 109.7 | | |
| PV1608 | Switzerland | Pully | 46.51 | 6.67 | 150 | 25.7 | X | |
| PV1610* | Switzerland | Pully | 46.51 | 6.67 | 150 | 27.2 | X | |
| PV1612 | Switzerland | Pully | 46.51 | 6.67 | 150 | 22.4 | X | |
| PV1873 | Switzerland | Cugnasco | 46.18 | 8.92 | 150 | 28.7 | X | |
| PV1933 | Switzerland | Pully | 46.51 | 6.67 | 150 | 25.3 | X | |
| PV1989 | France | Pouilly le Monial | 45.96 | 4.65 | 100 | 84.4 | | |
| PV2090 | France | Arles | 43.68 | 4.63 | 150 | 23.9 | X | |
| PV2282 | Spain | Sant Sadurni D'Anoia | 41.43 | 1.79 | 150 | 34.1 | X | |
| PV2416 | Italy | Pianello Val Tidone | 44.93 | 9.38 | 150 | 27.0 | | |
| PV2598 | Spain | Agoncillo | 42.47 | -2.29 | 150 | 31.1 | X | |
| PV2649 | France | Nogent-l'Abesse | 49.24 | 4.16 | 150 | 39.7 | X | |
| PV3174 | Italy | Piateda | 46.16 | 9.94 | 150 | 15.5 | X | |
| PV3175 | Italy | Canevino | 44.94 | 9.28 | 150 | 38.9 | | |
| PV3190 | Georgia | NA | NA | NA | 150 | 34.0 | | |
| PV3195 | Bulgaria | Vidin | 43.99 | 22.87 | 150 | 22.1 | X | |

Location: town or region where the individual was collected. Mean coverage: mean read coverage for the whole genome. Used for reference-free SNP analysis: Xs indicate that the individual has been used for the de novo SNP detection with discoSNP++.

*: testers used for mating-type phenotyping

†: resequencing was not successful for this individual

**Supplementary Table 2 Mating-type phenotyping for *Plasmopara viticola* individuals**
All individuals (rows) have been crossed with the six testers (columns). Successful crosses (i.e. oospores were observed after crossing) are indicated by black cells. Individuals successfully mating with P2 testers were determined as having the P1 mating type, and vice versa.

| | P1 testers | | | P2 testers | | | |
|---|---|---|---|---|---|---|---|
| | **PV413** | **PV1538** | **PV1610** | **PV1238** | **PV221** | **PV1400** | Mating type |
| PV13 | | | | ■ | ■ | ■ | P1 |
| PV125 | ■ | | | | | | P2 |
| **PV221** | ■ | ■ | | | | | **P2** |
| PV319 | | | | | ■ | | P1 |
| PV321 | ■ | ■ | | | | | P2 |
| PV330 | | | | ■ | ■ | ■ | P1 |
| PV334 | | | | | ■ | | P1 |
| PV336 | | | ■ | | | | P2 |
| PV340 | ■ | ■ | | | | | P2 |
| PV349 | | ■ | | | | | P2 |
| PV365 | | | ■ | | | | P2 |
| PV366 | ■ | | ■ | | | | P2 |
| PV368 | ■ | | | | | | P2 |
| PV375 | | | ■ | | | | P2 |
| PV382 | ■ | ■ | | | | | P2 |
| PV412 | | | | ■ | | | P1 |
| **PV413** | | | | ■ | ■ | | **P1** |
| PV414 | ■ | | | | | | P2 |
| PV1085 | | | | | ■ | | P1 |
| PV1094 | ■ | ■ | ■ | | | | P2 |
| PV1102 | ■ | ■ | ■ | | | | P2 |
| PV1130 | | | | ■ | | | P1 |
| PV1140 | ■ | ■ | ■ | | | | P2 |
| PV1174 | ■ | | | | | | P2 |
| PV1185 | | | | ■ | | | P1 |
| PV1194 | | | | ■ | | ■ | P1 |
| PV1225 | | | ■ | | | | P2 |
| **PV1238** | ■ | ■ | | | | | **P2** |
| PV1246 | ■ | ■ | ■ | | | | P2 |
| PV1272 | ■ | ■ | | | | | P2 |
| PV1286 | | | | ■ | ■ | ■ | P1 |
| PV1305 | | | ■ | | | | P2 |
| PV1314 | | ■ | | | | | P2 |
| PV1322 | | | | ■ | | | P1 |
| PV1346 | | | | ■ | ■ | | P1 |
| **PV1400** | ■ | ■ | | | | | **P2** |
| PV1456 | | | | ■ | | | P1 |
| PV1505 | | | | ■ | ■ | | P1 |
| **PV1538** | | | | ■ | ■ | | **P1** |
| PV1606 | | ■ | | | | | P2 |
| PV1608 | | | | | ■ | | P1 |
| **PV1610** | | | | ■ | ■ | | **P1** |
| PV1612 | ■ | ■ | | | | | P2 |
| PV1873 | | | | ■ | ■ | | P1 |
| PV1933 | | ■ | ■ | | | | P2 |
| PV1989 | | | | ■ | | ■ | P1 |
| PV2090 | | | | ■ | | | P1 |
| PV2282 | | | | ■ | | ■ | P1 |
| PV2416 | | ■ | | | | | P2 |
| PV2598 | ■ | ■ | ■ | | | | P2 |
| PV2649 | | ■ | | | | | P2 |
| PV3174 | | | | ■ | ■ | | P1 |
| PV3175 | | | | | ■ | | P1 |
| PV3190 | | | | ■ | | | P1 |
| PV3195 | | | | ■ | ■ | | P1 |

Individual IDs in bold designate tester individuals

**Supplementary table 3 Functional annotation of all candidate genes in the mating-type locus of *Plasmopara viticola***

| Protein ID | Length | Coverage TE (%) | Panther definition | GO terms | InterPro domains | KEGG definition | KEGG pathway | Non-syn mutations |
|---|---|---|---|---|---|---|---|---|
| PVIT_0008596.T1 | 135 | 0.00 | | | | | | |
| PVIT_0008599.T1 | 101 | 99.01 | | nucleic acid binding | | | | |
| PVIT_0008600.T1 | 795 | 16.60 | | transferase activity, transferring acyl groups; transferase activity | | | | |
| PVIT_0008601.T1 | 328 | 38.41 | | | | | | |
| PVIT_0008603.T1 | 193 | 79.27 | | | | | | |
| PVIT_0008604.T1 | 358 | 80.45 | FAMILY NOT NAMED | transferase activity, transferring acyl groups; transferase activity | Endonuclease/exonuclease/phosphatase | | | |
| PVIT_0008605.T1 | 752 | 25.66 | | transferase activity, transferring acyl groups; transferase activity | | | | |
| PVIT_0008608.T1 | 713 | 0.00 | | transferase activity, transferring acyl groups; transferase activity | | | | Ile234Thr, Tyr239Cys |
| PVIT_0008609.T1 | 462 | 82.47 | FAMILY NOT NAMED | DNA binding; transferase activity, transferring acyl groups; transferase activity | Endonuclease/exonuclease/phosphatase | | | |
| PVIT_0008615.T1 | 91 | 95.60 | | nucleic acid binding; zinc ion binding; metal ion binding; DNA integration | | | | |
| PVIT_0008616.T1 | 1375 | 0.00 | AAA-FAMILY ATPASE | protein targeting to peroxisome; peroxisome organization; peroxisomal membrane; peroxisome; protein binding; ATP binding; ATPase activity, coupled | ATPase, AAA-type, conserved site; UBX domain; AAA+ ATPase domain; ATPase, AAA-type, core; SEP domain; Peroxisome biogenesis factor 1, N-terminal, psi beta-barrel fold; Peroxisome biogenesis factor 1; P-loop containing nucleoside triphosphate hydrolase | PEX1; peroxin-1 | Peroxisome | Pro179Ser, Asp392Gly, Glu540Asp, Arg763Lys |
| PVIT_0008617.T1 | 160 | 0.00 | RECQ-MEDIATED GENOME INSTABILITY PROTEIN 2 | | | RMI2; RecQ-mediated genome instability protein 2 | Fanconi anemia pathway | |
| PVIT_0008618.T1 | 363 | 0.00 | PROTEIN DISULFIDE ISOMERASE | integral component of membrane; isomerase activity; cell redox homeostasis; protein folding; endoplasmic reticulum; protein disulfide isomerase activity; response to endoplasmic reticulum stress | Thioredoxin, conserved site; Disulphide isomerase; Thioredoxin-like fold; Thioredoxin domain | TXNDC5, ERP46; thioredoxin domain-containing protein 5 | Protein processing in endoplasmic reticulum | |
| PVIT_0008619.T1 | 83 | 0.00 | FAMILY NOT NAMED | Mo-molybdopterin cofactor biosynthetic process; cell part | Sulfur carrier ThiS/MoaD-like; Molybdopterin converting factor, subunit 1; Beta-grasp domain superfamily; Molybdopterin synthase/thiamin biosynthesis sulphur carrier, beta-grasp | MOCS2A, CNXG; molybdopterin synthase sulfur carrier subunit | Sulfur relay system | |
| PVIT_0008620.T1 | 303 | 50.50 | PEROXISOMAL MEMBRANE PROTEIN PEX13 | membrane; integral component of membrane | | PEX13; peroxin-13 | Peroxisome | Val79Gly, Leu233Pro, Leu233Phe, Glu237Asp |
| PVIT_0010914.T1 | 419 | 0.00 | SUBFAMILY NOT NAMED | integral component of membrane; transferase activity, transferring glycosyl groups | Glycosyltransferase, GlcNAc | | | |
| PVIT_0010915.T1 | 270 | 0.00 | FAMILY NOT NAMED | hydrolase activity, acting on carbon-nitrogen (but not peptide) bonds; nitrogen compound metabolic process | Carbon-nitrogen hydrolase | NTA1; protein N-terminal amidase [EC:3.5.1.-] | | |
| PVIT_0010916.T1 | 218 | 92.66 | SENTRIN/SUMO-SPECIFIC PROTEASE | proteolysis; cysteine-type peptidase activity | Ulp1 protease family, C-terminal catalytic domain | SENP8, NEDP1, DEN1; sentrin-specific protease 8 [EC:3.4.22.68] | | |
| PVIT_0010917.T1 | 449 | 0.00 | | DNA binding; cytoplasm; nucleus | Ubiquitin interacting motif; Proteasomal ubiquitin receptor Rpn13/ADRM1 | RPN13; 26S proteasome regulatory subunit N13 | Proteasome;Epstein-Barr virus infection | |
| PVIT_0010918.T1 | 765 | 0.00 | CATION EFFLUX PROTEIN/ ZINC TRANSPORTER | zinc ion transmembrane transporter activity; membrane; integral component of membrane; cation transmembrane transporter activity; cation transmembrane transport; Golgi apparatus; regulation of sequestering of zinc ion; cation transport; response to zinc ion; transmembrane transport | Cation efflux protein; Cation efflux transmembrane domain superfamily | SLC30A5_7, ZNT5_7, MTP, MSC2; solute carrier family 30 (zinc transporter), member 5/7 | | |
| PVIT_0010919.T1 | 137 | 0.00 | DYNEIN LIGHT CHAIN ROADBLOCK-TYPE 2 | microtubule-based movement; cytoplasmic dynein complex | Roadblock/LAMTOR2 domain | DYNLRB, DNCL2; dynein light chain roadblock-type | | |
| PVIT_0010920.T1 | 570 | 0.00 | | | | | | |
| PVIT_0010921.T1 | 84 | 0.00 | | | | | | |
| PVIT_0010922.T1 | 203 | 0.00 | HEPATOCELLULAR CARCINOMA-ASSOCIATED ANTIGEN | methyltransferase activity; transferase activity; methylation | Lysine methyltransferase | METTL21A; protein N-lysine methyltransferase METTL21A [EC:2.1.1.-] | | |
| PVIT_0010923.T1 | 265 | 0.00 | | | | | | |
| PVIT_0010924.T1 | 388 | 0.00 | | thiol-dependent ubiquitin-specific protease activity; proteolysis; protein binding | PDZ domain; Ankyrin repeat-containing domain; Ankyrin repeat | | | |

| | | | | | | | | |
|---|---|---|---|---|---|---|---|---|
| PVIT_0010925.T1 | 1071 | 10.92 | FYVE-FINGER-CONTAINING RAB5 EFFECTOR PROTEIN RABENOSYN-5-RELATED | protein binding; metal ion binding | FYVE zinc finger; WW domain; Zinc finger, FYVE-related; Zinc finger, FYVE/PHD-type; Zinc finger, RING/FYVE/PHD-type; START-like domain superfamily | | | |
| PVIT_0010926.T1 | 993 | 27.39 | | calcium ion binding | EF-Hand 1, calcium-binding site; EF-hand domain; EF-hand domain pair | | | |
| PVIT_0010927.T1 | 1230 | 0.00 | BACTERICIDAL PERMEABILITY-INCREASING BPI PROTEIN-RELATED | lipid binding; protein binding | PDZ domain; Bactericidal permeability-increasing protein, alpha/beta domain superfamily | | | |
| PVIT_0010928.T1 | 633 | 0.00 | DNA REPAIR POLYMERASE UMUC / TRANSFERASE FAMILY MEMBER | damaged DNA binding; transferase activity; DNA repair | UmuC domain; DNA polymerase, Y-family, little finger domain | POLH; DNA polymerase eta [EC:2.7.7.7] | Platinum drug resistance;Fanconi anemia pathway | His163Tyr |
| PVIT_0010929.T1 | 1479 | 0.00 | PATCHED-RELATED | lipid transporter activity; integral component of membrane; cell division; translational initiation; lipid transport; translation initiation factor activity | Sterol-sensing domain; Protein patched/dispatched | NPC1; Niemann-Pick C1 protein | Lysosome;Cholesterol metabolism | Ile297Met, Leu402Met, Glu1211Asp |
| PVIT_0010931.T1 | 417 | 68.35 | SERINE/ARGININE RICH SPLICING FACTOR | nucleic acid binding; nucleotide binding | RNA recognition motif domain; Nucleotide-binding alpha-beta plait domain superfamily | | | Asp229Gly, Leu284Arg, Ser294Gly, Ser303Gly, Thr306Ser, Gly310Ser, Gly325Val |
| PVIT_0010932.T1 | 390 | 77.69 | PHOSPHODIESTERASE HL | | ADP-ribosylation factor-like 2-binding protein, domain | | | Gly3Asp, Leu5Phe, Ser52Thr, His159Gln, Lys219Glu, premature start |
| PVIT_0010933.T1 | 479 | 33.19 | | nucleic acid binding; organic cyclic compound binding; nucleotide binding; cellular metabolic process; heterocyclic compound binding; intracellular; catalytic activity | HRDC domain; HRDC-like superfamily | | | Asp130Asn, Lys176Arg, Leu165Ile, Val249Ile, Thr250Ser |
| PVIT_0010934.T1 | 151 | 86.75 | | nucleic acid binding; DNA integration | | | | |
| PVIT_0010935.T1 | 161 | 60.87 | | nucleic acid binding; DNA binding; zinc ion binding; metal ion binding; DNA integration | | | | |
| PVIT_0010940.T1 | 76 | 75.00 | | | | | | |
| PVIT_0010941.T1 | 92 | 0.00 | | | | | | Gly82Cys |
| PVIT_0027771.T1 | 444 | 0.00 | MITOGEN-ACTIVATED PROTEIN KINASE | protein kinase activity; protein phosphorylation; nucleotide binding; ATP binding; phosphorylation | Protein kinase, ATP binding site; Protein kinase domain; Protein kinase-like domain superfamily | | | |
| PVIT_0027772.T1 | 842 | 0.00 | SPERMATOGENESIS-ASSOCIATED PROTEIN 5 | ATP binding | ATPase, AAA-type, conserved site; AAA+ ATPase domain; P-loop containing nucleoside triphosphate hydrolase | VCP, CDC48; transitional endoplasmic reticulum ATPase | Protein processing in endoplasmic reticulum;Legionellosis | |

Length: length of the protein sequence. Coverage TE: % of the protein sequence covered by a known transposable element sequence from the RepBase database. Non-syn mutations: Non-synonymous mutations caused by alternate alleles (i.e. present in the MAT-b allele) for SNPs associated with the mating-type phenotype.

**Supplementary table 4 Orthologs of the *Plasmopara viticola* mating type genes in the genomes of *Phytophthora infestans* , *Bremia lactucae* and *Plasmopara halstedii***

| *Plasmopara viticola* | | *Phytophthora infestans* | | *Bremia lactucae* | | *Plasmopara halstedii* | |
|---|---|---|---|---|---|---|---|
| Gene | Genomic coordinates | Gene | Genomic coordinates | Gene | Genomic coordinates | Gene | Genomic coordinates |
| PVIT_0008615.T1 | Plvit020 : 268538-268813 | - | - | - | - | Plhal710r1_S037g16879 | Plhal710r1_S037 : 514445-514675 |
| PVIT_0008616.T1 | Plvit020 : 284031-290118 | PITG_06660T0 | supercont1.8 : 3555056-3559199 | TDH71904.1 | SHOA01000004.1 : 3465462-3469694 | Plhal710r1_S058g22075 | Plhal710r1_S058 : 253289-257441 |
| PVIT_0008617.T1 | Plvit020 : 290201-290843 | PITG_06658T0 | supercont1.8 : 3553223-3553655 | - | - | Plhal710r1_S058g22076 | Plhal710r1_S058 : 257449-258211 |
| PVIT_0008618.T1 | Plvit020 : 290981-292072 | PITG_06657T0 | supercont1.8 : 3552016-3553107 | TDH71905.1 | SHOA01000004.1 : 3471347-3472438 | Plhal710r1_S058g22077 | Plhal710r1_S058 : 258213-259360 |
| PVIT_0008619.T1 | Plvit020 : 292484-292735 | - | - | TDH72680.1 | SHOA01000004.1 : 3312387-3312638 | Plhal710r1_S058g22080 | Plhal710r1_S058 : 270145-270396 |
| PVIT_0008620.T1 | Plvit020 : 292778-293875 | PITG_13927T0 | supercont1.30 : 1209885-1210997 | TDH72681.1 | SHOA01000004.1 : 3311222-3312707 | Plhal710r1_S058g22081 | Plhal710r1_S058 : 270418-271480 |
| PVIT_0010914.T1 | Plvit030 : 664127-665563 | PITG_19514T0 | supercont1.113 : 42045-43651 | TDH72556.1 | SHOA01000004.1 : 2470640-2472250 | Plhal710r1_S038g16979 | Plhal710r1_S038 : 178602-180207 |
| PVIT_0010915.T1 | Plvit030 : 666787-667975 | - | - | TDH72560.1 | SHOA01000004.1 : 2432641-2433906 | Plhal710r1_S038g16969 | Plhal710r1_S038 : 159428-160444 |
| PVIT_0010916.T1 | Plvit030 : 668239-668997 | PITG_18649T0 | supercont1.71 : 380428-381093 | TDH72113.1 | SHOA01000004.1 : 2674193-2674840 | Plhal710r1_S038g16968 | Plhal710r1_S038 : 158665-159334 |
| PVIT_0010917.T1 | Plvit030 : 669103-671428 | PITG_18650T0 | supercont1.71 : 381201-383324 | TDH72114.1 | SHOA01000004.1 : 2672938-2674102 | Plhal710r1_S038g16967 | Plhal710r1_S038 : 157440-158663 |
| PVIT_0010918.T1 | Plvit030 : 671952-674249 | PITG_18652T0 | supercont1.71 : 386886-388831 | TDH72111.1 | SHOA01000004.1 : 2668398-2670467 | Plhal710r1_S038g16964 | Plhal710r1_S038 : 151191-153602 |
| PVIT_0010919.T1 | Plvit030 : 674409-674822 | PITG_18659T0 | supercont1.71 : 509731-510138 | - | - | Plhal710r1_S038g16963 | Plhal710r1_S038 : 150556-150984 |
| PVIT_0010920.T1 | Plvit030 : 674890-677139 | PITG_18658T0 | supercont1.71 : 508361-509646 | - | - | Plhal710r1_S038g16962 | Plhal710r1_S038 : 148198-150509 |
| PVIT_0010922.T1 | Plvit030 : 678416-679027 | PITG_18657T0 | supercont1.71 : 505463-506107 | TDH72357.1 | SHOA01000004.1 : 3035742-3036617 | Plhal710r1_S038g16961 | Plhal710r1_S038 : 147238-147907 |
| PVIT_0010924.T1 | Plvit030 : 698965-700465 | PITG_18655T0 | supercont1.71 : 499107-500480 | TDH71914.1 | SHOA01000004.1 : 3038525-3042607 | - | - |
| PVIT_0010925.T1 | Plvit030 : 700575-703790 | PITG_18656T0 | supercont1.71 : 500643-504444 | TDH72358.1 | SHOA01000004.1 : 3038038-3041208 | Plhal710r1_S038g16945 | Plhal710r1_S038 : 117114-120572 |
| PVIT_0010926.T1 | Plvit030 : 711559-714862 | PITG_18644T0 | supercont1.71 : 200175-203646 | - | - | - | - |
| PVIT_0010927.T1 | Plvit030 : 715181-719081 | PITG_18641T0 | supercont1.71 : 143372-147280 | - | - | Plhal710r1_S058g22006 | Plhal710r1_S058 : 116854-120892 |
| PVIT_0010928.T1 | Plvit030 : 719171-721298 | PITG_18642T0 | supercont1.71 : 147405-149483 | TDH72635.1 | SHOA01000004.1 : 3068903-3071123 | Plhal710r1_S058g21970 | Plhal710r1_S058 : 58220-61000 |
| PVIT_0010929.T1 | Plvit030 : 722330-726769 | PITG_21540T0 | supercont1.436 : 41521-43864 | TDH72218.1 | SHOA01000004.1 : 3229267-3233712 | Plhal710r1_S058g21968 | Plhal710r1_S058 : 52165-56527 |
| PVIT_0010931.T1 | Plvit030 : 739521-740979 | PITG_06695T0 | supercont1.8 : 3784993-3786396 | TDH72050.1 | SHOA01000004.1 : 3516872-3518268 | Plhal710r1_S038g16929 | Plhal710r1_S038 : 79475-80755 |
| PVIT_0010932.T1 | Plvit030 : 740980-742486 | PITG_06694T0 | supercont1.8 : 3783665-3784865 | - | - | Plhal710r1_S038g16928 | Plhal710r1_S038 : 77507-79384 |
| PVIT_0010933.T1 | Plvit030 : 749107-750546 | PITG_06702T0 | supercont1.8 : 3915861-3917400 | TDH72557.1 | SHOA01000004.1 : 2434271-2435659 | Plhal710r1_S038g16937 | Plhal710r1_S038 : 98939-100506 |
| PVIT_0027771.T1 | Plvit600 : 27269-28831 | PITG_06701T0 | supercont1.8 : 3914139-3915718 | TDH72558.1 | SHOA01000004.1 : 2435764-2437367 | Plhal710r1_S038g16935 | Plhal710r1_S038 : 94334-95948 |
| PVIT_0027772.T1 | Plvit600 : 28900-31757 | PITG_06700T0 | supercont1.8 : 3912033-3914071 | - | - | Plhal710r1_S038g16936 | Plhal710r1_S038 : 95950-98826 |

Genomic coordinates are indicated as scaffold_ID : start_position-end_position, with 1-based coordinates.
